# Genomic prediction using machine learning: A comparison of the performance of regularized regression, ensemble, instance-based and deep learning methods on synthetic and empirical data

**DOI:** 10.1101/2022.06.09.495423

**Authors:** Vanda M. Lourenço, Joseph O. Ogutu, Rui A.P. Rodrigues, Hans-Peter Piepho

## Abstract

The accurate prediction of genomic breeding values is central to genomic selection in both plant and animal breeding studies. Genomic prediction involves the use of thousands of molecular markers spanning the entire genome and therefore requires methods able to efficiently handle high dimensional data. Not surprisingly, machine learning methods are becoming widely advocated for and used in genomic prediction studies. These methods encompass different groups of supervised and unsupervised learning methods. Although several studies have compared the predictive performances of individual methods, studies comparing the predictive performance of different groups of methods are rare. However, such studies are crucial for identifying (i) groups of methods with superior genomic predictive performance and assessing (ii) the merits and demerits of such groups of methods relative to each other and to the established classical methods. Here, we comparatively evaluate the genomic predictive performance and computational cost of several groups of supervised machine learning methods, specifically, *regularized regression* methods, *deep, ensemble* and *instance-based* learning algorithms, using one simulated animal breeding dataset and three empirical maize breeding datasets obtained from a commercial breeding program. Our results show that the relative predictive performance and computational expense of the groups of machine learning methods depend upon both the data and target traits and that for classical regularized methods, increasing model complexity can incur huge computational costs but does not necessarily always improve predictive accuracy. Thus, despite their greater complexity and computational burden, neither the adaptive nor the group regularized methods clearly improved upon the results of their simple regularized counterparts. This rules out selection of one procedure among machine learning methods for routine use in genomic prediction. The results also show that, because of their competitive predictive performance, computational efficiency, simplicity and therefore relatively few tuning parameters, the classical linear mixed model and regularized regression methods are likely to remain strong contenders for genomic prediction. The dependence of predictive performance and computational burden on target datasets and traits call for increasing investments in enhancing the computational efficiency of machine learning algorithms and computing resources.

**Author summary:** Machine learning methods are well suited for efficiently handling high dimensional data. Particularly, supervised machine learning methods have been successfully used in genomic prediction or genome-enabled selection. However, their comparative predictive accuracy is still poorly understood, yet this is a critical issue in plant and animal breeding studies given that increasing methodological complexity can substantially increase computational complexity or cost. Here, we show that predictive performance is both data and target trait dependent thus ruling out selection of one method for routine use in genomic prediction. We also show that for this reason, relatively low computational complexity and competitive predictive performance, the classical linear mixed model approach and regularized regression methods remain strong contenders for genomic prediction.

## Introduction

Rapid advances in genotyping and phenotyping technologies have enabled widespread and growing use of genomic prediction (GP). The very high dimensional nature of both genotypic and phenotypic data, however, is increasingly limiting the utility of the classical statistical methods. As a result, machine learning (ML) methods able to efficiently handle high dimensional data are becoming widely used in GP. It is therefore important to establish the relative predictive performance of different groups of ML methods. Even so, the predictive performance of groups of ML methods has attracted relatively little attention. The rising importance of ML methods in plant and animal breeding research and practice, increases both the urgency and importance of evaluating the relative predictive performance of groups of ML methods relative to each other and to classical methods. This can facilitate identification of groups of ML methods that balance high predictive accuracy with low computational cost for routine use with high dimensional phenotypic and genomic data, such as for GP, say.

ML is perhaps one of the most widely used branches of contemporary artificial intelligence. Using the ML methods facilitates automation of model building, learning and efficient and accurate predictions. The ML algorithms can be subdivided into two major classes: supervised and unsupervised learning algorithms. Supervised regression ML methods encompass regularized regression methods, deep, ensemble and instance-based learning algorithms. Supervised ML methods have been successfully used to predict genomic breeding values for unphenotyped genotypes, a crucial step in genome-enabled selection [25, 40, 41, 42, 43, 44, 45, 46, 49]. Furthermore, several studies have assessed the relative predictive performance of supervised ML methods in GP, including two ensemble methods and one instance-based method [44]; four regularized and two adaptive regularized methods [45]; three regularized and five regularized group methods [46] and several deep learning methods [40, 41, 42, 43, 49]. However, no study has comprehensively evaluated the comparative predictive performance of all these groups of methods relative to each other or to the classical regularized regression methods. We therefore rigorously evaluate the comparative predictive performance as well as the computational complexity or cost of three groups of popular and state-of-the-art ML methods for GP using one simulated animal dataset and three empirical datasets obtained from a commercial maize breeding program. We additionally offer brief overviews of the mathematical properties of the methods with emphasis on their salient properties, strengths and weaknesses and relationships with each other and with the classical regularization methods.

The rest of the paper is organized as follows. First we present the synthetic and real datasets. Second, we detail the methods compared in this study. Next, the results from the comparative analyses of the data are presented. Finally, a discussion of the results and closing remarks follow.

## Data

### Simulated (animal) data

We consider one simulated dataset [46], an animal breeding outbred population simulated for the 16-th QTLMAS Workshop 2012. The dataset consists of 4020 individuals genotyped for 9969 SNP markers. Out of these, 3000 individuals were phenotyped for three quantitative milk traits and the remaining 1020 were not phenotyped (see [46] for details). The goal of the analysis of the simulated dataset is to predict the genomic breeding values (PGBVs) for the 1020 unphenotyped individuals using the available genomic information. The simulated dataset also provides true genomic breeding values (TGBVs) for the 1020 genotypes for all the traits.

As in [46], to enable model fitting for the grouping methods, markers were grouped by assigning consecutive SNP markers systematically to groups of sizes 10, 20, …, 100 separately for each of the five chromosomes. Typically, the last group of each grouping scheme has fewer SNPs than the prescribed group size. Table 1 summarizes the simulated phenotypic data and highlights differences in the magnitudes of the three simulated quantitative traits *T*_1_, *T*_2_ and *T*_3_.

**Table 1.**
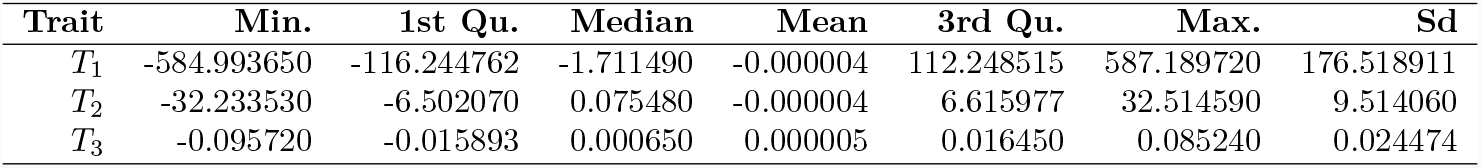
Summary statistics for the three quantitative traits (*T*_1_, *T*_2_ and *T*_3_) in the simulated training dataset (*n* = 3000 genotypes).

### Real (plant) data

For the application to empirical data sets, we use three empirical maize breeding datasets produced by KWS (breeding company) for the Synbreed project during 2010, 2011 and 2012. We first performed separate phenotypic analyses for each of the three real maize data sets to derive the adjusted means used in genomic prediction using a single stage mixed model assuming that genotypes are uncorrelated. The fixed effect in the mixed model comprised a tester (Tester) with two levels, genotypic group (GRP) with three levels, Tester × GRP and Tester × GRP×G (G=genotype). The random factors were location (LOC), trial (TRIAL) nested within location, replicate (REP) nested within trial and block (BLOCK) nested within replicate. The fitted random effects were LOC, LOC×TRIAL, LOC×TRIAL×REP, LOC×TRIAL×REP×BLOCK, Tester×GRP×SWITCH2×G and Tester×GRP×SWITCH1×G2. SWITCH1 and SWITCH2 in the last two effects are operators explained in greater detail in [13]. All the three maize datasets involved two testers and three genotypic groups. Accordingly, prior to genomic prediction, we accounted for and removed the effect of the tester×genotypic group (GRP) effect from the adjusted means (lsmeans) of maize yield by computing the arithmetic mean of the lsmeans for the interaction of testers with GRP for the genotyped lines. This mean was then subtracted from the lsmeans for each tester×GRP interaction term. The resulting deviations were subtracted from the lsmeans of the individual genotypes corresponding to each Tester×GRP interaction. This enabled us not to consider the Tester×GRP effect in the genomic prediction model. The SAS codes used for the preceding phenotypic analysis and computation of the adjusted genotype means used as the response variable in genomic prediction is provided in S5 File. For all the years, every line was genotyped for 32217 SNP markers. A subset of the SNP markers with non-zero variances were split into groups of sizes 10, 20, 30, 40, 50, 60, 70, 80, 90 and 100. Groups were defined by systematically grouping consecutive and spatially adjacent markers, separately for each of 10 chromosomes. The SAS code used to define the groups is provided in S6 File. All the checks (standard varieties) and check markers were deleted prior to genomic prediction. More details specific to the three datasets follow (Table 2 summarizes the number of genotypes in the training and validation datasets). The true breeding values are not known in this case.

**Table 2.** Number of genotypes in the training dataset (folds F1-F4) and validation dataset (fold F5) for each of the 10 cross-validation replicates for the 2010, 2011 and 2012 KWS real maize datasets.

|  | Folds | 2010 |  | 2011 |  | 2012 |  |
| --- | --- | --- | --- | --- | --- | --- | --- |
|  |  | F1-F4 | F5 | F1-F4 | F5 | F1-F4 | F5 |
| Training |  | 859 | 856 | 685 | 688 | 1104 | 1108 |
| Validation |  | 214 | 217 | 172 | 169 | 277 | 273 |

The **2010** dataset: the phenotypic dataset consists of 1073 individuals genotyped for 32217 SNPs and randomly split into 5 datasets, which we call folds, for 5-fold cross validation. The random splitting procedure was repeated 10 times to yield 10 replicates. In folds 1 − 4, 859 individuals (across all replicates) are used for training and 214 (across all replicates) individuals are used for validation. In fold 5, 856 individuals (across all replicates) are used for training and 217 (across all replicates) individuals are used for validation.

The **2011** dataset: the phenotypic dataset consists of 857 individuals genotyped for 32217 SNPs and randomly split into 5 datasets, which we call folds, for 5-fold cross validation. The random splitting procedure was repeated 10 times to yield 10 replicates. In folds 1 − 4, 685 individuals (across all replicates) are used for training and 172 (across all replicates) individuals are used for validation. In fold 5, 688 individuals (across all replicates) are used for training and 169 (across all replicates) individuals are used for validation.

The **2012** dataset: the phenotypic dataset consists of 1381 individuals genotyped for 32217 SNPs and randomly split into 5 datasets, which we call folds, for 5-fold cross validation. The random splitting procedure was repeated 10 times to yield 10 replicates. In folds 1 − 4, 1104 individuals (across all replicates) are used for training and 277 (across all replicates) individuals are used for validation. In fold 5, 1108 individuals (across all replicates) are used for training and 273 (across all replicates) individuals are used for validation.

Table 3 summarizes the KWS phenotypic data for 2010, 2011 and 2012. Each data split for each year (2010, 2011 and 2012) contained approximately 20% of the phenotypic observations and was obtained using stratified random sampling using the algorithm S7 File of [60] as modified in S8 File. The strata were defined by the combinations of the two testers and three genotypic groups.

**Table 3.**
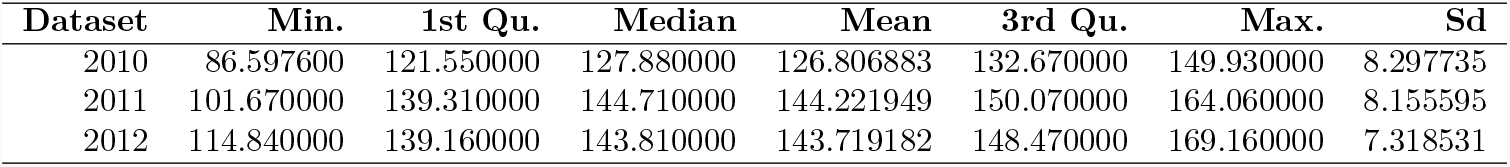
Summary statistics for maize yield in the KWS real maize datasets for 2010, 2011 and 2012.

## Methods

In this section we describe the four supervised ML groups of methods.

### Regularized regression methods

Consider the general linear regression model

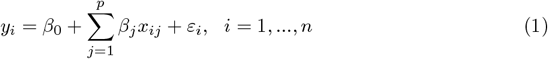

where *y*_*i*_ is the *i*-th observation of the response variable, *x*_*ij*_ is the *i*-th observation of the *j*-th covariate (*p* is the number of all covariates), *β*_*j*_ are the regression coefficients (unknown fixed parameters), *ε*_*i*_ are i.i.d. random error terms with *E*(*ε*_*i*_) = 0 and *var* 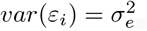, where 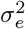 is an unknown random variance, and *n* is the sample size. The ordinary least squares estimator of ***β*** = (*β*_0_, …, *β*_*p*_)^′^, which is unbiased, is obtained by minimizing the residual sum of squares (RSS), i.e.,

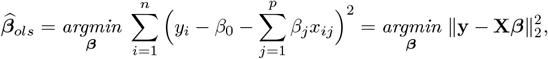

where

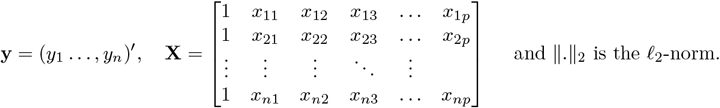

This estimator is typically not suitable when the design matrix **X** is less than full rank (**X** has a full rank if the number of its linearly independent rows or columns *k* = min(*p, n*)) or is close to collinearity (i.e., the covariates are close to being linear combinations of one another) [16]; problems that are frequently associated with *p >> n*. In genomic prediction (GP) one is interested in estimating the *p* regression coefficients *β*_*j*_ so that genomic breeding values of non-phenotyped genotypes can be predicted from the fitted model. The response variable **y** is often some quantitative trait and the *β*_*j*_’s are the coefficients of molecular markers spanning the whole genome, usually Single Nucleotide Polymorphisms (SNPs). Because in GP typically *p >> n*, the ordinary least squares (OLS) estimator breaks down and thus other methods for estimating ***β*** in (1) must be sought. Indeed, the increasingly high dimensional nature of high-throughput SNP-marker datasets has prompted increasing use of the power and versatility of regularization methods in genomic prediction to simultaneously select and estimate important markers and account for multicollinearity [44, 45].

Without loss of generality, we assume, consistent with the standard practice in regularized estimation where a distance-based metric is used for prediction, that the response variable is mean-centered whereas the covariates in (1) are standardized, so that

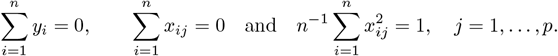

Regularized regression methods minimize a non-negative loss function (RSS or other) plus a non-negative penalty function. Standardizing the covariates prior to model fitting ensures that the penalty is applied evenly to all covariates. Mean-centering the response and the covariates is usually done for notational simplicity but also eliminates the need to estimate the intercept *β*_0_.

After the penalized models have been fit, the final estimates are obtained by back transformation to the original scale by re-introducing an intercept (*β*_0_). In particular, for a mean-centered response **y** and standardized predictor **X**^∗^, predictions are obtained by

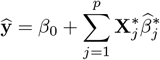

with 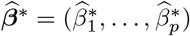, the regression coefficients from the model fit with the mean-centered response **y** and standardized covariates 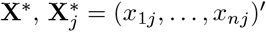 the *j*-th covariate and 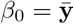. One can also choose to predict using the original predictor **X**^∗^ without standardization. In that case one should back transform the 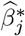 to the original scale and consider

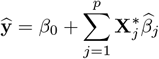

with 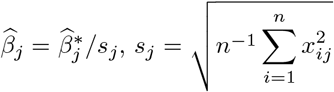 the standard deviation of the j-th covariate 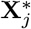 and 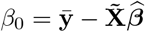, where 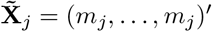 is a vector of size *n* with *m*_*j*_ being the mean of the *j*-th covariate 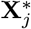.

The primary goal of regularization methods is to reduce model complexity resulting from high dimensionality by reducing the number of predictors in the model. This is achieved by either shrinking some coefficients to become exactly zero, and so drop out of the model, or shrinking all coefficients to be close to zero and each other but not exactly zero. Ideally, a desirable estimator of ***β*** should (i) correctly select the nonzero coefficients with probability converging to 1 (i.e. with near certainty; *selection consistency*) and (ii) yield estimators of the nonzero coefficients that are asymptotically normal with the same means and covariances that they would have if the zero coefficients were known exactly in advance (*asymptotic normality*). An estimator satisfying these two conditions is said to possess the *oracle* property [14, 15].

For the remainder of the paper, we assume that **X** is a *n* × *p* marker matrix (e.g., with the genotypes {*aa, Aa, AA*} coded as {0, 1, 2} or {−1, 0, 1} for *p* biallelic SNPs under an additive model) with **X**_*j*_ denoting the *j*-th SNP covariate and ***β*** = (*β*_1_, …, *β*_*p*_) denoting the unknown vector of marker effects. Table 4 (upper half) summarizes the methods discussed in this sub-section.

**Table 4.** A summary of the estimators and penalty functions for the bridge-type and adaptive bridge-type regularized regression methods used in this study. The adaptive methods have an *a* prefix in their names.

| Method | Penalty | Estimator |  |
| --- | --- | --- | --- |
| bridge | $p_{\lambda,\gamma}(\boldsymbol{\beta}) = \lambda \sum_{j=1}^p \vert \beta_j\vert ^\gamma$ | $\hat{\boldsymbol{\beta}}_{bridge} = \underset{\boldsymbol{\beta}}{\operatorname{argmin}} \left\{ \text{RSS} + \lambda \sum_{j=1}^p \vert \beta_j\vert ^\gamma \right\}, \gamma > 0, \lambda \geq 0$ | (2) |
| • $\gamma = 1$: | | | |
| LASSO | $p_\lambda(\boldsymbol{\beta}) = \lambda \ \boldsymbol{\beta}\ _1$ | $\hat{\boldsymbol{\beta}}_{lasso} = \underset{\boldsymbol{\beta}}{\operatorname{argmin}} \left\{ \text{RSS} + \lambda \ \boldsymbol{\beta}\ _1 \right\}$ | (3) |
| • $\gamma = 2$: | | | |
| ridge | $p_\lambda(\boldsymbol{\beta}) = \lambda \ \boldsymbol{\beta}\ _2^2$ | $\hat{\boldsymbol{\beta}}_{ridge} = \underset{\boldsymbol{\beta}}{\operatorname{argmin}} \left\{ \text{RSS} + \lambda \ \boldsymbol{\beta}\ _2^2 \right\}$ | (4) |
| • Combination of LASSO and ridge penalties ($\gamma = 1, 2$, respectively): | | | |
| ENET | $p_\lambda(\boldsymbol{\beta}) = \lambda_1 \ \boldsymbol{\beta}\ _1 + \lambda_2 \ \boldsymbol{\beta}\ _2^2$ | $\hat{\boldsymbol{\beta}}_{enet} = (1 + \lambda_2) \times \left\{ \underset{\boldsymbol{\beta}}{\operatorname{argmin}} \text{RSS} + \lambda_1 \ \boldsymbol{\beta}\ _1 + \lambda_2 \ \boldsymbol{\beta}\ _2^2 \right\}$ | (5) |
| abridge | $p_{\lambda,\gamma}(\boldsymbol{\beta}) = \lambda \sum_{j=1}^p w_j \vert \beta_j\vert ^\gamma$ | $\hat{\boldsymbol{\beta}}_{abridge} = \underset{\boldsymbol{\beta}}{\operatorname{argmin}} \left\{ \text{RSS} + \lambda \sum_{j=1}^p w_j \vert \beta_j\vert ^\gamma \right\}$ | (6) |
| • $\gamma = 1$: | | | |
| aLASSO | $p_\lambda(\boldsymbol{\beta}) = \lambda \ \mathbf{w}\boldsymbol{\beta}\ _1$ | $\hat{\boldsymbol{\beta}}_{alasso} = \underset{\boldsymbol{\beta}}{\operatorname{argmin}} \left\{ \text{RSS} + \lambda \ \mathbf{w}\boldsymbol{\beta}\ _1 \right\}$ | (7) |
| • Combination of aLASSO and ridge penalties ($\gamma = 1, 2$, respectively): | | | |
| aENET | $p_\lambda(\boldsymbol{\beta}) = \lambda_1 \ \mathbf{w}\boldsymbol{\beta}\ _1 + \lambda_2 \ \boldsymbol{\beta}\ _2^2$ | $\hat{\boldsymbol{\beta}}_{aenet} = k \times \left( \underset{\boldsymbol{\beta}}{\operatorname{argmin}} \left\{ \text{RSS} + \lambda_1 \ \mathbf{w}\boldsymbol{\beta}\ _1 + \lambda_2 \ \boldsymbol{\beta}\ _2^2 \right\} \right)$ | (8) |

### Bridge-type estimators

The most popular regularization methods in genomic prediction include ridge regression (RR; [26]), the least absolute shrinkage and selection operator (LASSO; [57]) and the elastic net (ENET; [67]). All these methods are special cases of the bridge estimator [16, 20] given by

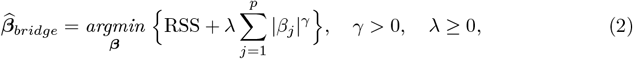

where the *regularization* parameter λ balances the goodness-of-fit against model complexity and the *shrinkage* parameter *γ* determines the order of the penalty function. The optimal combination of λ and *γ* can be selected adaptively for each dataset by grid search using cross-validation (CV; if the focus is on predictive performance) or by information criteria (e.g., AIC or BIC; if the focus is on model fit). Bridge regression automatically selects relevant predictors when 0 *< γ* ≤ 1, shrinks the coefficients when *γ >* 1 and reduces to subset selection when *γ* = 0. The bridge estimator reduces to the LASSO estimator when *γ* = 1 and to the ridge estimator when *γ* = 2. Specifically,

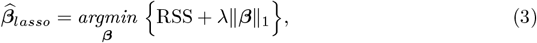

where ∥.∥_1_ is the *ℓ*_1_-norm, and

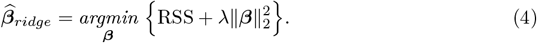

The bridge estimator also enjoys several other useful and interesting properties (see [28, 33] for more details). We summarize these salient properties with emphasis on the special cases of the LASSO (*γ* = 1) and the ridge estimators (*γ* = 2).

1. The asymptotic properties of bridge estimators have been studied by [28]. In particular, for *p < n* with *p* increasing to infinity with *n* and under appropriate regularity conditions, bridge estimators enjoy the oracle property for 0 *< γ <* 1, implying that, neither the LASSO nor the ridge estimator enjoys the oracle property [14, 15]. If *p >> n* and no assumptions are imposed on the covariate matrix, then the regression parameters are generally non identifiable. However, if a suitable structure is assumed for the covariate matrix, then bridge estimators achieve consistent variable selection and estimation [28].
2. Although the LASSO estimator performs automatic variable selection, it is a biased and inconsistent estimator [64, 65]. Moreover, it is unstable with high-dimensional data because it
  i. cannot select a larger number of predictors *p* than the sample size *n* if *p >> n*;
  ii. arbitrarily selects one member of a set of pairwise highly correlated predictors and ignores the other.
3. The ridge estimator performs well for many predictors each of which has a small effect but cannot shrink the coefficients to become exactly zero. Moreover, the ridge estimator
  i. prevents coefficients of linear regression models with many correlated variables from being poorly determined and exhibiting high variance;
  ii. shrinks coefficients of correlated predictors equally towards zero and towards each other;
  iii. retains all predictor variables in the model leading to complex and less interpretable models.
  In addition, RR has close connections with marker-based BLUP and genomic BLUP [38], which we clarify in what follows. The ridge estimator is given by

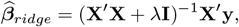

where, if *λ* is estimated by cross-validation as suggested above, then the ridge estimator may be denoted by RR-CV. Another way of looking at the ridge estimator is to assume in (1) that 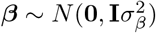 is a random vector of unknown marker effects and that 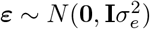 is an unknown random error term, where 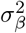 and 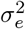 are the unknown marker-effect and error variances, respectively. This model is now a linear mixed model and hence, the variances can be estimated via the restricted maximum likelihood (REML) method. The BLUP solution for the marker effects under model (1) is given by

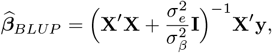

establishing the equivalence of BLUP and RR [51, 54] and that one can actually estimate the ridge parameter *λ* by 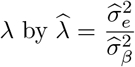. Because we use REML to estimate the two variance components in 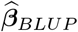, we refer to this RR appproach as RR-REML. The underlying mixed model for gBLUP (ignoring fixed effects and considering just one random effect per individual) is

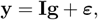

where, 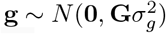 is the random vector of unknown genotypic (breeding) values, 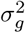 is the unknown genetic variance, ***ε*** and 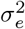 are defined as before, and **G** = **XX**^′^ is the genomic (marker-based) relationship matrix. The genomic-estimated breeding values (GEBVs; 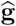) are equivalent to the estimates 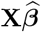 from model (1) [24, 50]. Indeed, for this special case 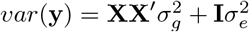 which is exactly the same as that for model (1) when ***β*** is taken as random.
4. Due to the nature of the *ℓ*_1_ penalty, particularly for high values of *λ*, the LASSO estimator will shrink many coefficients to exactly zero, something that never happens with the ridge estimator.

#### Elastic net estimator

The elastic net estimator blends two bridge-type estimators, the LASSO and the ridge, to produce a composite estimator that reduces to the LASSO when *λ*_2_ = 0 and to the ridge when *λ*_1_ = 0. Specifically, the elastic net estimator is specified by

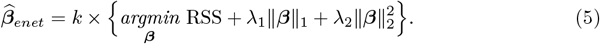

with *k* = 1 + *λ*_2_ if the predictors are standardized (as we assume) or *k* = 1 + *λ*_2_*/n* otherwise. Even when *λ*_1_, *λ*_2_ ≠ 0, the elastic net estimator behaves much like the LASSO but with the added advantage of being robust to extreme correlations among predictors. Moreover, the elastic net estimator is able to select more than *n* predictors when *p >> n*. Model sparsity occurs as a consequence of the *ℓ*_1_ penalty term. Mazumder et al. [36] proposed an estimation procedure based on sparse principal components analysis (PCA), which produces an even more sparse model than the original formulation of the elastic net estimator [67]. Because it blends two bridge-type estimators, neither of which enjoys the oracle property, the ENET also lacks the oracle property.

Other competitive regularization methods that are asymptotically oracle efficient (*p < n* with *p* increasing to infinity with *n*), which do not fall into the category of bridge-type estimators, are the *smoothly clipped absolute deviations* (SCAD [15, 31]) and the *minimax concave penalty* (MCP [63, 65]) methods. Details of the penalty functions and other important properties of both methods can be found elsewhere [5, 46].

### Adaptive regularized regression methods

The adaptive regularization methods are extensions of the regularized regression methods that allow the resulting estimators to achieve the oracle property under certain regularity conditions. Table 4 (lower half) summarizes the adaptive methods considered here.

#### Adaptive bridge-type estimators

Adaptive bridge estimators extend the bridge estimators by introducing weights in the penalty term. More precisely,

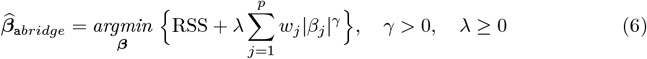

Where 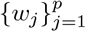 are adaptive data-driven weights. As with the bridge-type estimator, the adaptive bridge estimator simplifies to the adaptive LASSO (aLASSO) estimator when *γ* = 1 and to the adaptive ridge estimator when *γ* = 2. Chen et al. [10] studied the properties of adaptive bridge estimators for the particular case when *p < n* (with *p* increasing to infinity with *n*), 0 *< γ <* 2 and 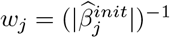 with 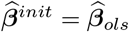. They showed that for 0 *< γ <* 1, and under additional model assumptions, adaptive bridge estimators enjoy the oracle property. For 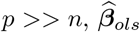 cannot be computed and thus other initial estimates, such as 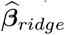, have to be used. Theoretical properties of the adaptive bridge estimator for *p >> n* do not seem to have been well studied thus far. The adaptive LASSO estimator was proposed by [68] to remedy the problem of the lack of the oracle property of the LASSO estimator [14, 15]. The penalty for the adaptive LASSO is given by (adaptive bridge estimator with *γ* = 1)

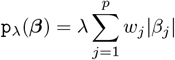

where the adaptive data-driven weights 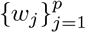 can be computed as 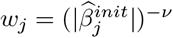 with 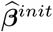 an initial root-*n* consistent estimate of ***β*** obtained through least squares (or ridge regression if multicollinearity is important) and *v* is a positive constant. Consequently,

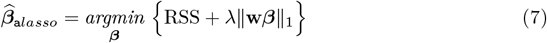

with *v* chosen appropriately, performs as well as the oracle, i.e., the adaptive LASSO achieves the oracle property. Nevertheless, this estimator still inherits the LASSO’s instability with high dimensional data. The values of *λ* and *v* can be simultaneously selected from a grid of values, with values of *v* selected from {0.5, 1, 2}, using two-dimensional cross-validation [68].

Grandvalet [21] shows that the adaptive ridge estimator (adaptive bridge estimator with *γ* = 2) is equivalent to the LASSO in the sense that both produce the same estimate and thus the adaptive ridge is not considered further.

#### Adaptive elastic-net

The adaptive elastic-net (aENET) combines the ridge and aLASSO penalties to achieve the oracle property [70] while at the same time alleviating the instability of the aLASSO with high dimensional data. The method first computes 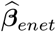 as described above for the elastic net estimator, then constructs the adaptive weights as 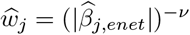, where *v* is a positive constant, and then solves

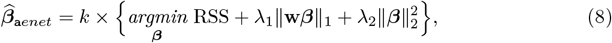

where *k* = 1 + *λ*_2_ if the predictors are standardized (as we assume) or *k* = 1 + *λ*_2_*/n* otherwise. In particular, when *λ*_2_ = 0 the adaptive elastic-net reduces to the aLASSO estimator. This is also the case when the design matrix is orthogonal regardless of the value of *λ*_2_ [67, 68, 70].

Other adaptive regularization methods are the *multi-step adaptive ENET* (maENET), the *adaptive smoothly clipped absolute deviations* (aSCAD) and the *adaptive minimax concave penalty* (aMCP) methods. Details of the penalty functions and noteworthy properties of the latter three methods are summarized elsewhere [45, 59].

### Regularized group regression methods

Regularized regression methods that select individual predictors do not exploit information on potential grouping structure among markers, such as that arising from the association of markers with particular Quantitative Trait Loci (QTL) on a chromosome or haplotype blocks, to enhance the accuracy of genomic prediction. The nearby SNP markers in such groups are linked, producing highly correlated predictors. If such grouping structure is present but is ignored by using models that select individual predictors only, then such models may be inefficient or even inappropriate, reducing the accuracy of genomic prediction [46]. Regularized group regression methods are regularized regression methods with penalty functions that enable the selection of the important groups of covariates and include group bridge (`g`bridge), group LASSO (`g`LASSO), group SCAD (`g`SCAD) and group MCP (gMCP) methods (see [1, 6, 30, 46, 47, 61] for detailed reviews). Some grouping methods such as the group bridge, sparse group LASSO (`sg`LASSO) and group MCP, besides allowing for group selection, also select the important members of each group [4] and are therefore said to perform bi-level selection, i.e., group-wise and within-group variable selection. Bi-level selection is appropriate if predictors are not distinct but have a common underlying grouping structure.

Estimators and penalty functions for the regularized grouped methods can be formulated as follows. Consider subsets *A*_1_, …, *A*_*L*_ of {1, …, *p*} (*L* being the total number of covariate groups), representing known covariate groupings of design vectors, which may or may not overlap. Let 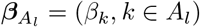 be the regression coefficients in the *l*-th group and *p*_*l*_ the cardinality of the *l*-th group (i.e., the number of unique elements in *A*_*l*_). Regularized group regression methods estimate 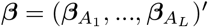 by minimizing

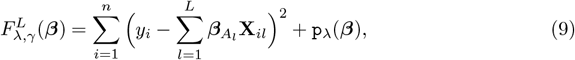

where **X**_.*l*_ is a matrix with columns corresponding to the predictors in group *l*.

Because 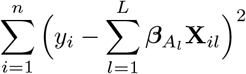 in (9) is equivalent to RSS some authors use the RSS formulation directly. It is assumed that all the covariates belong to at least one of the groups. Table 5 summarizes the methods described in this section.

**Table 5.** Penalty functions and estimators for some group regularized regression methods used in this study.

| Method | Penalty | Estimator |
| --- | --- | --- |
| gbridge | $p_{\lambda, \gamma}(\beta) = \lambda \sum_{l=1}^L c_l \ \beta_{A_l}\ _1^{\gamma}$ | $\hat{\beta}_{\text{gbridge}} = \underset{\beta}{\operatorname{argmin}} \left\{ \text{RSS} + \lambda \sum_{l=1}^L c_l \ \beta_{A_l}\ _1^{\gamma} \right\} \quad (10)$ |
| gLASSO | $p_{\lambda}(\beta) = \lambda \sum_{l=1}^L \sqrt{p_l} \ \beta_{A_l}\ _2$ | $\hat{\beta}_{\text{glasso}} = \underset{\beta}{\operatorname{argmin}} \left\{ \text{RSS} + \lambda \sum_{l=1}^L \sqrt{p_l} \ \beta_{A_l}\ _2 \right\} \quad (11)$ |
| sgLASSO | $p_{\lambda, \alpha}(\beta) = \alpha \lambda \ \beta\ _1 + (1 - \alpha) \lambda \sum_{l=1}^L \sqrt{g_l} \ \beta_l\ _2$ | $\hat{\beta}_{\text{sglasso}} = \underset{\beta}{\operatorname{argmin}} \left\{ \text{RSS} + \alpha \lambda \ \beta\ _1 + (1 - \alpha) \lambda \sum_{l=1}^L \sqrt{g_l} \ \beta_l\ _2 \right\} \quad (12)$ |

### Group bridge-type estimators

Group bridge-type estimators use in (9) the penalty term 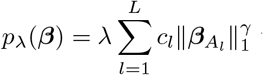 with *c*_*l*_ constants that adjust for the different sizes of the groups. The group bridge-type estimators are thus obtained as

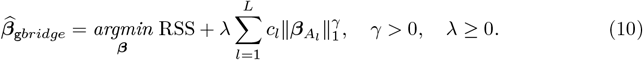

A simple and usual choice for the *c*_*l*_ constants consists in considering each 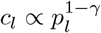. When 0 *< γ <* 1 group bridge can be used simultaneously for group and individual variable selection. Also, note that under these assumptions, the group bridge estimator correctly selects groups with nonzero coefficients with probability converging to one under reasonable regularity conditions, i.e., it enjoys the *oracle group selection* property (see [27] for details). When the group sizes are all equal to one, i.e., *p*_*l*_ = 1 ∀ 1≤ *l*≤ *L*, then group bridge estimators reduce to the bridge estimators.

#### Group LASSO and sparse group LASSO

Group LASSO regression uses in (9) the penalty function 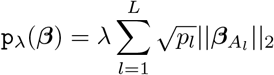. The group LASSO estimator is thus given by

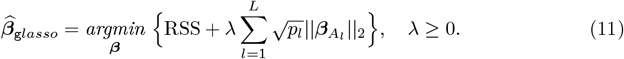

Unlike the group bridge estimator (0 *< γ <* 1), `g`LASSO is designed for group selection, but does not select individual variables within the groups. Indeed, its formulation is more akin to that of the adaptive ridge estimator [27]. As with the group-bridge estimator, when the group sizes are all equal to one, i.e., *p*_*l*_ = 1 ∀ 1 ≤ *l* ≤ *L*, the `g`LASSO estimator reduces to the LASSO estimator.

Because the `g`LASSO does not yield sparsity within a group (it either discards or retains a whole group of covariates) the sparse group lasso (`sg`LASSO), which blends the LASSO and the `g`LASSO penalties, was proposed [18, 56]. Specifically, the sgLASSO estimator is given by

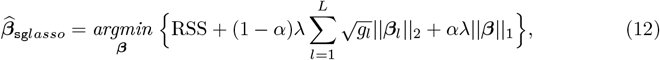

where *α* ∈ [0, 1] provides a convex combination of the lasso and group lasso penalties (*α* = 0 gives the `g`LASSO fit, *α* = 1 gives the LASSO fit). The `g`LASSO is superior to the standard LASSO under the strong group sparsity and certain other conditions, including a group sparse eigenvalue condition [29]. Because the `sgLASSO` lacks the oracle property, the adaptive sparse group LASSO was recently proposed to remedy this drawback [52].

Note that there are two types of sparsity, i.e., (i) “groupwise sparsity”, which refers to the number of groups with at least one nonzero coefficient, and (ii) “within group sparsity” that refers to the number of nonzero coefficients within each nonzero group. The “overall sparsity” usually refers to the total number of non-zero coefficients regardless of grouping.

Other group regularization methods are the *hierarchical group LASSO* (hLASSO), the *group smoothly clipped absolute deviations* (gSCAD) and the *group minimax concave penalty* (gMCP) methods. Details of the penalty functions and salient properties of these methods can be found in [35, 46, 48, 66**?**].

### Ensemble methods

Ensemble methods build multiple models using a given learning algorithm and then combine their predictions to produce an optimal estimate. The two most commonly used algorithms are *bagging* (or bragging) and *boosting*. Whereas *bagging* is a stagewise procedure that combines the predictions of multiple models (e.g., classification or regression trees) to yield an average prediction, *boosting* is a stagewise process in which each stage attempts to improve the predictions at the previous stage by up-weighting poorly predicted values. Below, we briefly discuss two popular ensemble methods, namely, random forests, an extension of bagging, and gradient boosting algorithms. Note that, although variable scaling (centering or standardizing) might accelerate convergence of the learning algorithms, the ensemble methods do not require it. Indeed, the collection of partition rules used with the ensemble methods should not change with scaling.

#### Random forests (RF)

The random forests algorithm is an ensemble algorithm that uses an ensemble of unpruned decision (classification or regression) trees, each grown using a bootstrap sample of the training data, and randomly selected (without replacement) subsets of the predictor variables (features) as candidates for splitting tree nodes. The randomness introduced by bootstrapping and selecting a random subset of the predictors reduces the variance of the random forest estimator, often at the cost of a slight increase in bias. The RF regression prediction for a new observation *y*, say 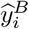, is made by averaging the output of the ensemble of B trees {*T*(*y*_*i*_, Ψ_*b*_}_*b*=1,…,*B*_ as [23]

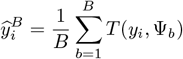

where Ψ_*b*_ characterizes the *b*-th RF tree in terms of split variables, cut points at each node, and terminal node values. Recommendations on how to select the number of trees to grow, the number of covariates to be randomly chosen at each tree node and the minimum size of terminal nodes of trees, below which no split is attempted, are provided by [9, 34]. We refer to [9, 23, 34] for further details on the RF regression.

#### Stochastic Gradient Boosting (SGB)

Boosting enhances the predictive performance of base learners such as classification or regression trees, each of which performs only slightly better than random guessing, to become arbitrarily strong [23]. As with RF, boosting algorithms can also handle interactions, nonlinear relationships, automatically select variables and are robust to outliers, missing data and numerous correlated and irrelevant variables. In regression, boosting is an additive expansion of the form

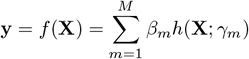

where *β*_1_, …, *β*_*M*_ are the expansion coefficients and the basis functions *h*(**X**; *γ*), base learners, are functions of the multivariate argument **X**, characterized by a set of parameters *γ* = (*γ*_1_, …, *γ*_*M*_). Typically these models are fit by minimizing a loss function *L* (e.g., the squared-error loss) averaged over the training data

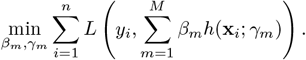

We used regression trees as basis functions in which the parameters *γ*_*m*_ are the splitting variables, split points at the internal nodes, and the predictions at the terminal nodes. Boosting regression trees involves generating a sequence of trees, each grown on the residuals of the previous tree. Prediction is accomplished by weighting the ensemble outputs of all the regression trees. We refer to [23, 55] for further details on SGB (see, e.g., [55] for the interpretation of boosting in terms of regression for a continuous, normally distributed response variable).

### Instance-based methods

For the instance-based methods, scaling before applying the method is crucially important. Scaling the variables (features) prior to model fitting prevents possible numerical difficulties in the intermediate calculations and helps avoid domination of numeric variables with smaller by those with greater magnitude and range.

#### Support Vector Machines

Support vector machines (SVM) is a popular supervised learning technique for classification and regression of a quantitative response *y* on a set of predictors, in which case the method is called support vector regression or SVR [58]. In particular, SVR uses the model

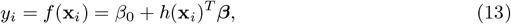

with **x**_*i*_ = (*x*_*i*1_, …, *x*_*ip*_)^′^ and where the approximating function *f* (**x**_*i*_) is a linear combination of basis functions *h*(**x**_*i*_)^*T*^, which can be linear (or nonlinear) transformations of **x**_*i*_. The goal of SVR is to find a function *f* such that *f* (**x**_*i*_) deviates from *y*_*i*_ by a value no greater than *ε* for each training point **x**_*i*_, and at the same time is as flat as possible. This so-called *ε*-insensitive SVR, or simply *ε*-SVR, thus fits a model (13) using only those residuals which are smaller in absolute value than *ε* and a linear loss function for larger residuals. The choice of the loss function (e.g., linear, quadratic, Huber) usually considers the noise distribution pertaining to the data samples, level of sparsity and computational complexity.

If Eq (13) is the usual linear regression model, i.e., 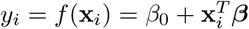, one considers the following minimization problem

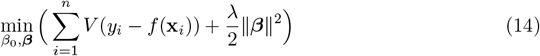

where *λ* is the regularization parameter (cost) that controls the trade-off between flatness and error tolerance, ∥.∥ refers to the norm under a Hilbert space (i.e., 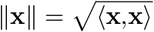 with **x** a *p* ≥ 1 dimensional vector) and

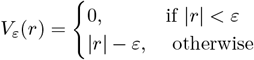

is an *ε*-insensitive linear loss. Given the minimizers of (14) 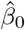 and 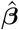, the solution function has the form

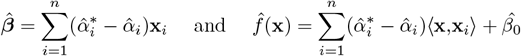

where 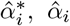 are positive weights given to each observation (i.e., to the column vector **x**_*i*_) estimated from the data. Typically only a subset of 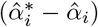 are non-zero with the observations associated to these so called *support vectors*, and thus the name of the method, SVM. More details on SVM can be found in [23].

### Deep learning methods

Deep learning (DL) algorithms are a special class of neural networks and encompass an assortment of architectures (e.g., convolutional, recurrent and densely connected neural networks) that depend on many parameters (hyperparameters) whose careful optimization is crucial to enhancing predictive accuracy and minimizing overfitting (see [2, 12, 39, 49, 62, 71] for further insights into DL architectures and other particulars and the supplementary materials https://github.com/miguelperezenciso/DLpipeline of [49] for a list of the main DL hyperparameters, their role and related optimization issues). It can be very challenging to achieve great improvements in predictive accuracy in genomic prediction studies with DL because hyperparameter optimization can be extremely demanding and also because DL requires very large training datasets which might not always be available [40, 41, 42, 43].

Regardless of the selected DL architecture and the adopted hyperparameter calibration, when building the neural network model an optimization method must also be selected. The three top ranked optimizers for neural networks are mini-batch gradient descent, gradient descent with momentum and adaptive moment estimation (ADAM; [32]). Among the three, the mini-batch gradient descent and Adam are usually preferred, because they perform well most of the time. In terms of convergence speed ADAM is often clearly the winner and thus a natural choice [53].

Next, we offer a few more details on the feed-forward neural network (FFNN), also known in the literature as a multi-layer perceptron (MLP), which, besides being one of the most popular DL architectures, is well suited for regression problems.

#### Feed-forward neural network (FFNN)

A feed-forward neural network (FFNN), is a neural network that does not assume a specific structure in the input features (i.e., in the covariates). This neural network consists of an input layer, an output layer and multiple hidden layers between the input and output layers.

The model for a FFNN with one hidden layer expressed as a multiple linear regression model (1) is given by

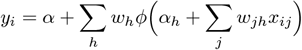

where the *y*_*i*_ (output) and *x*_*ij*_ (input) are defined as in model (1), *α* is the output bias, *h* runs over the units of the hidden layer, *α*_*h*_ refers to the bias of the *h*-th unit of the hidden layer, *w*_*jh*_ refer to the weights between the inputs and the hidden layer, *w*_*h*_ refer to the weights between the hidden layer and the output, *ϕ* is the *activation* function of the hidden layer. The model parameters *α, α*_*h*_, *w*_*h*_ and *w*_*jh*_ are unknown network parameters that need to be estimated by minimizing some fitting criterion such as least squares or some measure of entropy from some training data (network training). This model can be represented graphically as a set of inputs linked through a hidden layer to the outputs as shown in Fig 1.

**Fig. 1.**
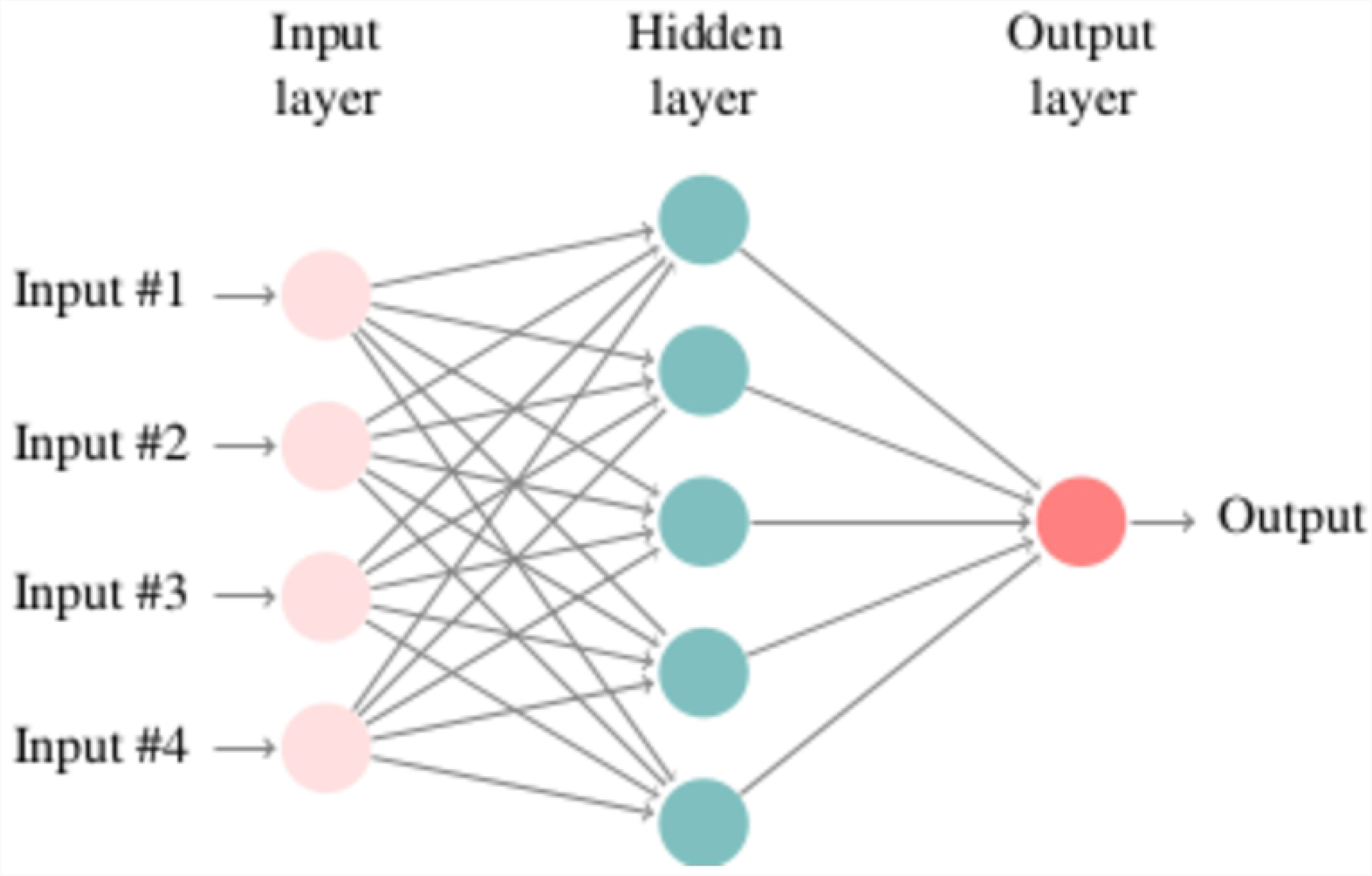
Graphical representation of a feed-forward neural network (FFNN) with one hidden layer.

Further details on neural networks in general and the FFNN in particular can be found in [23, 40, 41, 42, 43, 49]. Note that, to avoid potential numerical difficulties, it is recommended that both the target (response variable; here assumed to be continuous and normally distributed), and the features (covariates) are standardized prior to training the network [49].

### Performance assessment & noteworthy details of model fitting

#### Performance assessment

For the simulated dataset, we assessed predictive performance using predictive accuracy (PA), the Pearson correlation between the PGBVs and the TGBVs breeding values. For all the three KWS empirical data sets, predictive performance was expressed as predictive ability (PA), the Pearson correlation between the PGBVs and the observed genomic breeding values (OGBVs), also calculated using cross validation. The higher the PA, the better is the relative predictive performance of a method. We additionally assessed the predictive performance of the methods using the out-of-sample mean squared prediction error (MSPE) and the mean absolute prediction error (MAPE). Specifically,

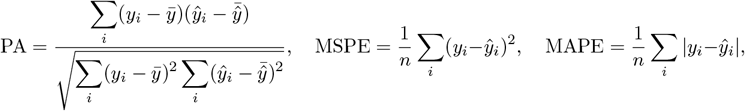

where the *y*_*i*_ and 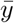 are, respectively, the TGBVs and mean TGBVs for the single simulated dataset, but the OGBVs and mean OGBVs for the empirical datasets, and the ŷ*_i_* are the PGBVs. 10-fold CV is used to assess the PA for each method for the simulated datasets in contrast to the 5-fold CV used with the three empirical maize datasets.

For the cross validation, we aimed to have at least 150 individuals per fold. Accordingly, each phenotypic dataset was randomly split into *k* approximately equal parts. The breeding values for each of the *k* folds were predicted by training the model on the *k* 1 remaining folds and a CV error (CVE) computed for each of the *k* folds. The method with the smallest CVE was selected to predict the breeding values for the unphenotyped genotypes for the simulated dataset, and the phenotyped genotypes in the validation sets for each of the three empirical maize datasets.

#### Noteworthy details of model fitting

Because we used CV for model selection, we fixed the data split whenever possible by setting a specific positive seed before splitting the data to enhance the reproducibility of results. Besides the *k*-fold split, common to all the methods, some methods involve additional kinds of randomization, making it impossible to reproduce results even if the data split or the seed for the random number generator are fixed prior to fitting the model (RF, SVM and FFNN methods). This happens because some of the routines internally generate some random number seeds when they initialize the computations. Consequently, this introduces an additional source of variation between results obtained from different runs or even from using different computing platforms. Ideally, for such cases, and at the risk of vastly increasing the computational burden, the model could be fitted a large number of times until the average and the range (or standard error) of PA stabilize. Even so, we considered only single model runs for the RF and SVM methods. For the FFNN method we implemented 1000 runs for the simulated data and 10 for the KWS data.

In addition, the following details of model fitting are noteworthy:

i. *γ* = 0.5 was considered in the bridge and grouped bridge methods.
ii. Calibration of random forests involves selecting the number of trees to grow (*n*_*trees*_), the random number of covariates to select for growing each tree (*n*_*covars*_) and the minimum size of the terminal nodes per tree (*n*_*nodes*_), below which no split is attempted. The parameters that affect the final accuracy the most are the first two. Increasing the *n*_*trees*_ only increases the accuracy up to some point but can substantially increase the computational time. Here, we fix *n*_*trees*_ = 1000 (this ensures that every input row gets predicted at least a couple of times) and *n*_*nodes*_ = 1 and search for the best value of *n*_*covars*_ in {0.5, 1, 2} × (*p/*3).
iii. For the SGB method we assumed the Gaussian distribution for minimizing squared-error loss. The basic boosting algorithm requires the specification of two parameters: the number of splits (*J*; or the number of nodes, which equals the number of splits plus one) and the number of trees (or iterations; *n*_*trees*_) to be fitted. Hastie et al [23] point out that the number of splits *J* such that 4 ≤ *J* ≤ 8 generally works well with results being fairly insensitive to particular choices in this range. We use *J* = 6. As for *n*_*trees*_, it should neither be too small (bad fit) nor too large (overfit). Usually, the suitable *n*_*trees*_ can be found by checking how well the model fits a test dataset, where a typical fraction of the data used for testing is 0.5 (can be much smaller if the dataset is very large). We search for *n*_*trees*_ in {500, 1500, 3000}. As with RF, we fix the number of terminal nodes per tree *n*_*nodes*_ = 1. In addition, we set the shrinkage factor applied to each tree in the expansion to 0.001 and the subsampling fraction to 0.5 [17].
iv. For the SVR method, we considered an insensititvity zone of *ε* = 0.001 across traits with the regularization parameter *λ* (cost) determined by grid-search over the values {0.01, 0.1, 1, 10, 100}.
v. We used the Adam optimizer, ‘Relu’ (rectified linear units) activation function and a linear output layer in the configuration of the FFNN.

Because we used the Python software and GPU to fit the neural networks, we were able to produce 1000 different runs of the FFNNs for the simulated dataset and 10 runs of the FFNNs for each of the three real datasets (amounting to 10 *runs* × 5 *folds* × 10 *reps* FFNNs fits for each real dataset), and report the average plus the range for the PA.

All the methods are implemented in the R software and are available in various R packages. Table 6 lists the R packages we used to analyse the synthetic and real datasets. For the FFNN method, and because of fine tuning requirements, we used the Python software and packages Numpy, Pandas and Tensorflow.

**Table 6.** List of R packages and routines used in this paper

| Type | Method | Package | Routine |
| --- | --- | --- | --- |
| Regularized | Bridge | grpreg [7] | cv.grpreg() |
|  | RR-CV, ENET | glmnet [19] | cv.glmnet() * |
|  | RR-REML | rrBLUP [11] | mixed.solve() |
|  | LASSO, SCAD, MCP | ncvreg [8] | cv.ncvreg() * |
| Sparse regularized | ENET | sparsenet [36] | cv.sparsenet() |
| Adaptive regularized | LASSO | glmnet [19] | cv.glmnet() * |
|  | SCAD |  | asnet() * |
|  | MCP | msaenet [59] | amnet() * |
|  | ENET |  | aenet() *, msaenet() * |
| Group regularized | Hierarchical LASSO | glinternet [35] | glinternet.cv() |
|  | Sparse LASSO | SGL [56] | cvSGL() |
|  | Bridge |  |  |
|  | LASSO, MCP, SCAD | grpreg [7] | cv.grpreg() |
|  | gel, cMCP (bi-level) |  |  |
| Ensemble | Random Forests | randomForest [9] | randomForest() |
|  | Stochastic Gradient Boosting | gbm [22] | gbm.fit() |
| Instance-based | Support Vector Machine | e1071 [37] | tune.svm() |

\* These functions allow for internal parallelization of computations.

## Results

Although we did not fully quantify the computational costs of the different methods, the computational burden increased strikingly from the simple regularized through the adaptive to the grouped methods. A similar trend was also apparent from the ensemble, through the instance-based to the deep learning methods. Computational time may be reduced greatly by parallelizing the estimation or optimization algorithms, but this strategy may not always be available and can be challenging to implement for some methods.

### Simulated (animal) data

The relative performances of the various methods on the simulated data varied with the target trait and with whether performance was assessed in terms of predictive accuracy or prediction error. Performance also varied in terms of computational cost with some methods requiring considerably more time than others. Results of genomic prediction accuracy for the simulated data are displayed in Tables 7 to 11. Table 12 reports the calibration details for the fitted FFNN.

**Table 7.**
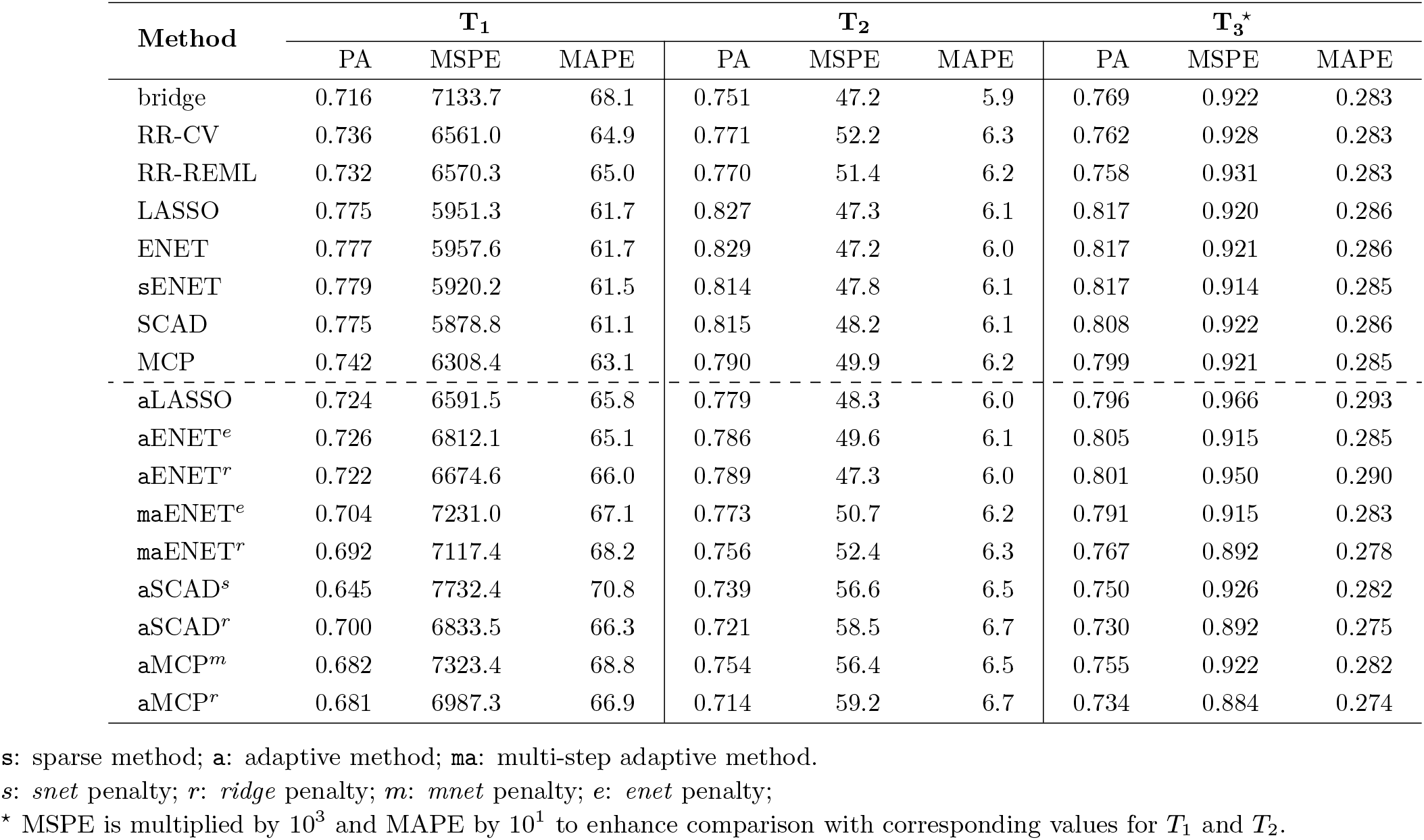
Prediction accuracy (PA) of the **regularized** and **adaptive regularized** methods, computed as the Pearson correlation coefficient between the true breeding values (TBVs) and the predicted breeding values (PBVs), for the simulated dataset, where *T*_1_ − *T*_3_ refer to three quantitative milk traits. The choice of *λ*, where applicable, was based on the 10-fold CV. The mean squared and absolute prediction errors are also provided.

Table 9 displays the range of the observed predictive accuracies across all the classes of the regularized methods for traits *T*_1_ −*T*_3_. Neither the adaptive nor the group regularized methods seem to improve upon the results of their regularized counterparts, although group regularized methods do provide some slight improvement upon the results of the adaptive regularized methods. Even though all the regularized regression methods had comparable overall performance, the best compromise between high PA (≥ 0.77 for *T*_1_, 0.82 for *T*_2_ and 0.81 for *T*_3_) and small prediction errors was achieved by the LASSO, ENET, sENET and SCAD (Table 7; first half). Within the class of adaptive regularized methods, the best compromise was achieved by aLASSO and aENET (Table 7; second half; PA ≥ 0.72 for *T*_1_, 0.78 for *T*_2_ and 0.80 for for *T*_3_). For the group regularized methods, a good compromise was achieved by the gLASSO and gSCAD (Table 8; mean PA ≥ values 0.76 for *T*_1_, 0.82 for *T*_2_ and 0.81 for *T*_3_). Whereas the worst performing group regularized methods in terms of the estimated PAs were the cMCP and gel for *T*_1_ (PA*<* 0.7), sgLASSO and gel for *T*_2_ (PA*<* 0.8) and hLASSO and gel for *T*_3_ (PA*<* 0.8), the worst performing methods in terms of prediction errors were the gel (*T*_1_ & *T*_2_ only) and sgLASSO (*T*_3_ only). Of all the group regularized methods, the most time consuming were the sgLASSO and hLASSO, with sgLASSO requiring several more months to compute results for trait *T*_1_ than for traits *T*_2_ or *T*_3_.

**Table 8.**
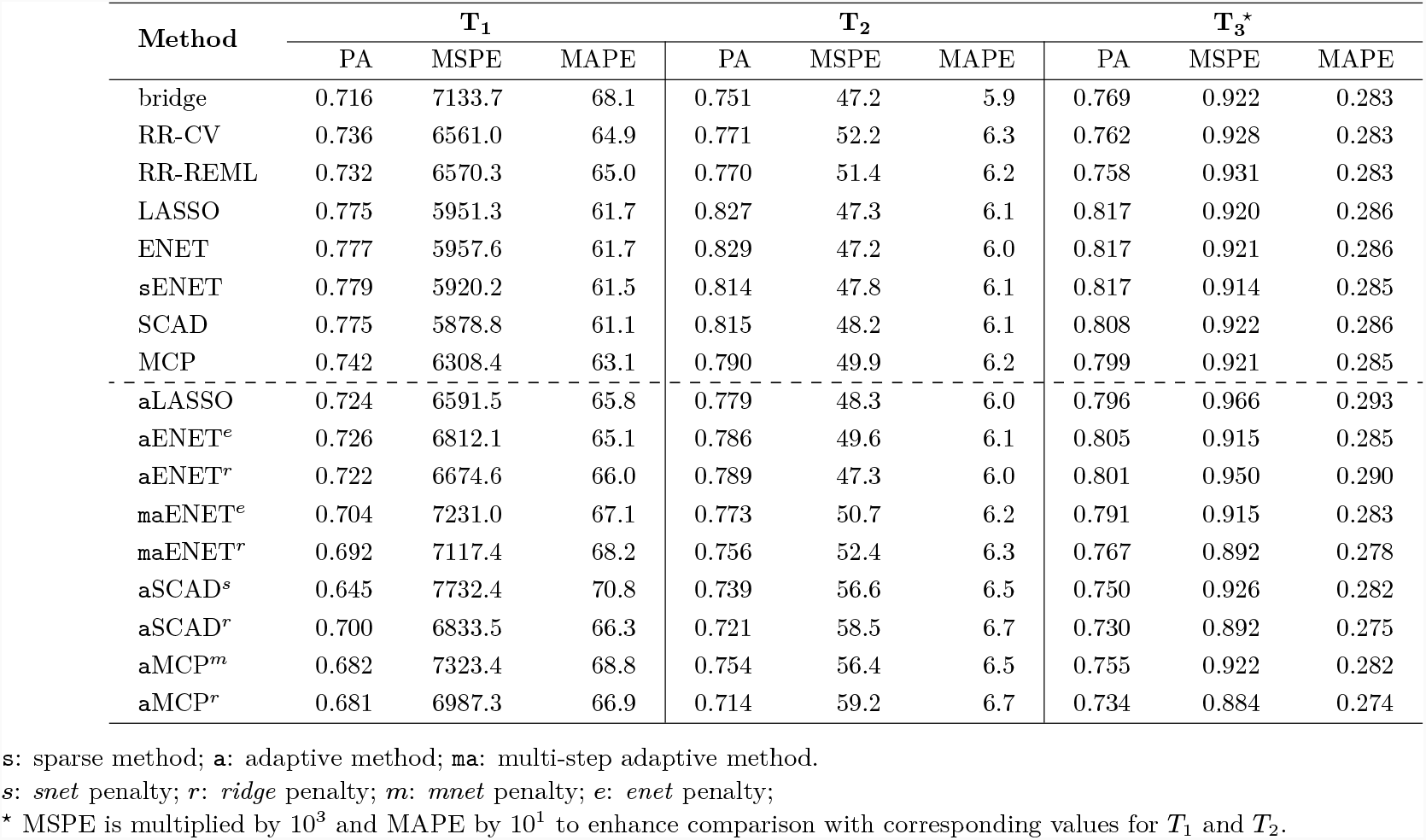
Prediction accuracy (PA) of the **group regularized** methods (mean and range values of PA across the different groupings), computed as the Pearson correlation coefficient between the true breeding values (TBVs) and the predicted breeding values (PBVs), for the simulated dataset, where *T*_1_ − *T*_3_ refer to three quantitative milk traits. Choice of *λ* was based on the 10-fold CV. Display refers to the mean, max and min values of PA across all the 10 grouping schemes. The mean squared and absolute prediction errors are also provided.

**Table 9.** Range of the estimated predictive accuracies across the classes of regularized methods for traits *T*_1_ − *T*_3_.

| | $T_1$ | $T_2$ | $T_3$ |
| --- | --- | --- | --- |
| Regularized | 0.716 – 0.779 | 0.770 – 0.829 | 0.758 – 0.817 |
| Adaptive Regularized | 0.645 – 0.726 | 0.714 – 0.789 | 0.730 – 0.805 |
| Group Regularized<sup>†</sup> | 0.653 – 0.766 | 0.758 – 0.820 | 0.765 – 0.814 |

<sup>†</sup> Values refer to the range of the observed mean PAs.

The ensemble, instance-based and deep learning methods did not improve upon the results of the regularized or the group regularized methods (Tables 10 & 11). Among the latter groups of methods, RF provided the best compromise between high PA and small prediction errors.

**Table 10.** Prediction accuracy (PA) of the **ensemble and instance-based** methods, computed as the Pearson correlation coefficient between the true breeding values (TBVs) and the predicted breeding values (PBVs), for the simulated dataset, where *T*_1_ − *T*_3_ refer to three quantitative milk traits.

| Method | $T_1$ | | | $T_2$ | | | $T_3^*$ | | |
| --- | --- | --- | --- | --- | --- | --- | --- | --- | --- |
|  | PA | MSPE | MAPE | PA | MSPE | MAPE | PA | MSPE | MAPE |
| Random Forests (RF)<sup>†</sup> | 0.741 | 7945.8 | 72.4 | 0.788 | 58.6 | 6.5 | 0.713 | 0.924 | 0.272 |
| Stochastic Gradient Boosting (SGB)* | 0.690 | 8382.5 | 73.8 | 0.725 | 67.6 | 7.0 | 0.676 | 0.950 | 0.275 |
| Support Vector Machines (SVM)<sup>‡</sup> | 0.695 | 7102.1 | 67.2 | 0.740 | 53.9 | 6.4 | 0.731 | 0.934 | 0.282 |

<sup>†</sup> reported values refer to a single run of the random forest with the subset of markers selected randomly for growing each tree set equal to $0.5 \times \frac{p}{3}$ ;
\* the best number of trees grown was 3000 for all traits;
<sup>‡</sup> reported values refer to a single run of the SVM with the best cost $\lambda = 100$ for trait 1 and $\lambda = 10$ for traits 2 and 3;
\* MSPE is multiplied by $10^3$ and MAPE by $10^1$ to enhance comparison with corresponding values for $T_1$ and $T_2$ .

**Table 11.** Prediction accuracy (PA) of the **deep learning** methods, computed as the Pearson correlation coefficient between the true breeding values (TBVs) and the predicted breeding values (PBVs), for the simulated dataset, where *T*_1_ − *T*_3_ refer to three quantitative milk traits.

| | | $T_1$ | | $T_2$ | | $T_3^*$ | |
| --- | --- | --- | --- | --- | --- | --- | --- |
|  |  | Mean | Range | Mean | Range | Mean | Range |
| Feed Forward Neural Networks (FFNN)<sup>†</sup> | PA | 0.705 | 0.692–0.717 | 0.725 | 0.705–0.741 | 0.703 | 0.651–0.731 |
|  | MSPE | 8092.5 | 6046.5–13556.7 | 60.6 | 32.5–102.4 | 0.974 | 0.464–1.790 |
|  | MAPE | 72.1 | 61.6–95.4 | 6.6 | 4.6–9.1 | 0.284 | 0.183–0.402 |

<sup>†</sup> values refer to 1000 different runs of the neural network;
\* MSPE is multiplied by $10^3$ and MAPE by $10^1$ to enhance comparison with corresponding values for $T_1$ and $T_2$ .

**Table 12.** Best FFNN model calibration parameters selected for each of the three quantitative milk traits *T*_1_ − *T*_3_.

| FFNN | T<sub>1</sub> | T<sub>2</sub> | T<sub>3</sub> |
| --- | --- | --- | --- |
| Number of hidden layers | 1 | 3 | 3 |
| Number of units (hidden layer) | 200 | 800 | 400 |
| Number of epochs | 200 | 260 | 300 |
| Dropout rate (input layer) | 0.1 | 0.15 | 0.15 |
| Batch size | 128 | 32 | 16 |
| Learning rate | 10<sup>-4</sup> | 2 × 10<sup>-5</sup> | 2 × 10<sup>-5</sup> |
| Dropout rate (hidden layers) | 0.8 | 0.8 | 0.5 |
| Dropout rate (last hidden layer) | 0.8 | 0.88 | 0.775 |
| Batch normalization (hidden layer) | No | Yes | Yes |

Predictive performance varied not only among the methods but also with the target quantitative traits. Specifically, trait *T*_3_ had the highest predictive accuracies for the adaptive methods, whereas trait *T*_2_ was generally top ranked across all the remaining methods.

### Real (plant) data

The ridge regression methods plus the overall best performing methods (high PA values and low prediction errors) for each class of methods based on the analysis of the simulated dataset, were applied to each of the three KWS empirical maize datasets. The specific methods fitted to the KWS maize datasets comprised RR-CV, RR-REML, sENET, aENET (enet penalty), gLASSO, RF and FFNN.

Results are displayed in Table 13. Across the three real maize datasets, the highest predicitive abilities were obtained for the 2010 dataset. The estimated predictive abilities (PA) are under 0.7 for the 2010 dataset but under 0.6 for the 2011 dataset and 2012 (except for RR-REML with an estimated PA of 0.616), regardless of the method used. RR-REML (2011 & 2012 datasets) and RF (2010 dataset) are evidently the best performing methods (higher PA values and lower prediction errors). On the other hand, aENET^*e*^ (2010 & 2011 datasets) and RF (2012 dataset) are the worst performing methods (lower PA and higher prediction errors). Interestingly, the RF performed both the best (2010 dataset) and the worst (2012 dataset), clearly emphasizing that the methods are strongly data dependent.

**Table 13.** Predictive ability (PA; mean and range values computed across the 5-fold validation datasets and 10 replicates) of the **regularized** and **adaptive regularized** methods, computed as the Pearson correlation coefficient between the observed breeding values (OBVs) and the predicted breeding values (PBVs), for the KWS datasets. The choice of *λ*, where applicable, was based on 4-fold CV.

| Method | 2010 |  |  | 2011 |  | 2012 |  |
| --- | --- | --- | --- | --- | --- | --- | --- |
|  |  | Mean | Range | Mean | Range | Mean | Range |
| RR-CV | PA | 0.632 | 0.529-0.724 | 0.555 | 0.441-0.648 | 0.598 | 0.549-0.674 |
|  | MSPE | 41.9 | 31.7-58.3 | 46.4 | 35.5-62.1 | 34.5 | 29.1-41.5 |
|  | MAPE | 4.7 | 4.1-5.2 | 5.2 | 4.6-6.1 | 4.6 | 4.3-5.0 |
| RR-REML | PA | 0.649 | 0.523-0.725 | 0.576 | 0.469-0.663 | 0.616 | 0.555-0.678 |
|  | MSPE | 41.2 | 31.2-59.2 | 44.4 | 35.4-63.4 | 33.4 | 27.8-40.2 |
|  | MAPE | 4.6 | 4.0-5.2 | 5.0 | 4.5-6.0 | 4.5 | 4.1-4.9 |
| sENET | PA | 0.626 | 0.502-0.723 | 0.527 | 0.372-0.639 | 0.596 | 0.531-0.697 |
|  | MSPE | 42.4 | 32.6-62.4 | 48.8 | 37.2-84.5 | 34.7 | 28.0-42.2 |
|  | MAPE | 4.7 | 4.3-5.3 | 5.3 | 4.6-6.8 | 4.6 | 4.2-5.1 |
| aENET<sup>e</sup> | PA | 0.592 | 0.453-0.701 | 0.482 | 0.326-0.651 | 0.572 | 0.501-0.676 |
|  | MSPE | 46.5 | 37.4-67.6 | 54.3 | 37.7-98.9 | 37.0 | 30.6-46.3 |
|  | MAPE | 4.9 | 4.5-5.6 | 5.6 | 4.9-7.2 | 4.7 | 4.4-5.1 |
| gLASSO<sup>‡</sup> | PA | 0.584 | 0.494-0.673 | 0.512 | 0.404-0.606 | 0.586 | 0.514-0.679 |
|  | MSPE | 46.3 | 36.0-59.8 | 49.5 | 37.5-65.6 | 35.5 | 28.1-42.5 |
|  | MAPE | 4.9 | 4.3-5.5 | 5.4 | 4.6-6.3 | 4.7 | 4.2-5.1 |
| Random Forests<sup>†</sup> | PA | 0.656 | 0.549-0.727 | 0.570 | 0.474-0.659 | 0.556 | 0.484-0.633 |
|  | MSPE | 39.7 | 28.9-56.3 | 45.7 | 34.1-61.4 | 37.3 | 29.9-46.3 |
|  | MAPE | 4.6 | 4.0-5.1 | 5.1 | 4.4-6.0 | 4.8 | 4.3-5.3 |
| Feed Forward Neural Networks<sup>*</sup> | PA | 0.634 | 0.514-0.728 | 0.541 | 0.412-0.656 | 0.586 | 0.512-0.663 |
|  | MSPE | 42.9 | 31.9-61.7 | 49.6 | 38.4-75.1 | 36.5 | 29.0-48.5 |
|  | MAPE | 4.7 | 4.0-5.4 | 5.3 | 4.6-6.6 | 4.7 | 4.2-5.3 |

**a:** adaptive method; **s:** sparse method; **g:** grouped method; *e:* *enet* penalty; <sup>‡</sup>reported values refer to the grouping indexes (or sizes) 50, 30 and 80 for the 2010, 2011 & 2012 datasets, respectively; <sup>†</sup>reported values refer to a single run of the random forest with the subset of markers selected randomly for growing each tree set equal to $0.5 \times \frac{p}{3}$ for the 2010 and 2011 datasets but to $2 \times \frac{p}{3}$ for the 2012 dataset; <sup>\*</sup> The best FFNN performing model, in terms of PA, for the 2010 and 2011 datasets was the one used for trait $T_3$ from the simulated data, whereas for the 2012 dataset, it was the one used for trait $T_1$ from the simulated data.

## Discussion

We have investigated the predictive performance of several state-of-the art machine learning methods in genomic prediction via the use of one simulated and three real datasets. All the methods showed reasonably high predictive performance for most practical selection decisions. But the relative predictive performance of the methods was both data and target trait dependent, complicating and precluding omnibus comparative evaluations of the genomic prediction methods, thus ruling out selection of one procedure for routine use in genomic prediction. If reproducibility of results, low computational cost and time are important considerations, then using the regularized regression methods comes highly recommended because they consistently produced, with relatively lower computational cost and computing time, reasonably accurate and competitive predictions relative to the other groups of methods for the simulated and the three real datasets. Even among the regularized regression methods, increasing model complexity from simple through the adaptive to grouped methods, generally only increased computing time without clearly improving predictive performance.

The ensemble, instance-based and deep-learning ML methods need the tuning of numerous hyperparameters thus requiring considerable computing time to adequately explore the entire hyperparameter space. This will not always be possible in most applications because of limiting time and computational resources leading to potentially less than optimal results and may well partly explain why these methods did not clearly outperform the classical ML methods. Indeed, the computational costs of the ensemble, instance-based and deep learning methods can quickly become prohibitive, if all the parameters are tuned by searching over the often large grid of values. This will typically require not only proficiency in programming and algorithm parallelization and optimization, but excellent computing resources. These constraints, plus the growing size of phenotypic and genomic data, make it difficult to identify methods for routine use in genomic prediction and call for greater focus on and investment in enhancing the computational efficiencies of algorithms and computing resources.

## Supporting information

**S1 File. S1 Data**. Simulated (animal breeding) dataset. (ZIP)

**S2 File. S2 Data**. Simulated dataset that mimics the original KWS real dataset set for 2010.

(ZIP)

**S3 File. S1 Code**. R codes used to fit the machine learning algorithms to the simulated (animal breeding) dataset.

(ZIP)

**S4 File. S2 Code**. Python code used to fit the deep learning (FFNN) algorithm to the simulated (animal breeding) dataset.

(ZIP)

**S5 File. S1 Text**. SAS code used for phenotypic analysis of the KWS real maize dataset and computation of the adjusted genotypic means used as the response variable in genomic prediction.

(DOCX)

**S6 File. S2 Text**. SAS code used to assign consecutive and spatially adjacent SNP markers on the same chromosome to groups of sizes 10, 20, 30, …, 100 for use with the grouped regularized regression models.

(DOCX)

**S7 File. S3 Text**. SAS Macro written by [60] used to split each of the KWS 2010, 2011 and 2012 datasets into 5 distinct parts based on a specified probability vector. (DOCX)

**S8 File. S4 Text**. SAS macro used to split each of the KWS 2010, 2011 and 2012 datasets into 5 distinct parts using stratified random sampling and the macro of [60].

(DOCX)

## Acknowledgements

We thank KWS for providing the maize datasets.

## Data availability

The simulated animal data from the QTLMAS workshop is provided in S1 File in the supplementary materials together with the R and Python codes used to analyse these data (respectively, S3 File and S4 File). The KWS data is proprietary data and cannot be shared publicly for confidentiality reasons. As a result, we provide a synthetic dataset that mimics the KWS data in S2 File, which can be used with our codes to illustrate the implementation of the machine learning methods.

## Author details

^1^ Center for Mathematics and Applications (CMA), FCT NOVA and Department of Mathematics, FCT NOVA, 2829–516 Caparica, Portugal. ^2^ Institute of Crop Science, Biostatistics Unit, University of Hohenheim, Fruwirthstrasse 23, 70599 Stuttgart, Germany.

## References

1. Bach, F. (2008). Consistency of the group lasso and multiple kernel learning. Journal of Machine Learning, 9, 1179–1225.

2. Bengio, Y. (2012). Practical recommendations for gradient-based training of deep architectures. In Neural Networks: Tricks of the trade (pp. 437–478). Springer, Berlin, Heidelberg.

3. Bien, J., Taylor, J. & Tibshirani, R. (2013). A lasso for hierarchical interactions. The Annals of Statistics, 41, 1111–1141.

4. Breheny, P. & Huang, J. (2009). Penalized methods for bi-level variable selection. Statistics Interface, 2, 369–380.

5. Breheny, P. & Huang, J. (2011). Coordinate descent algorithms for nonconvex penalized regression, with applications to biological feature selection. Annals of Applied Statistics, 5, 232–253.

6. Breheny, P., & Huang, J. (2015). Group descent algorithms for nonconvex penalized linear and logistic regression models with grouped predictors. Statistics and Computing, 25(2), 173–187.

7. Breheny, P. & Breheny, M. P. (2021). Package ‘grpreg’.

8. Breheny, P. & Breheny, M. P. (2021). Package ‘ncvreg’.

9. Breiman, L. (2001). Random forests. Machine Learning, 45, 5–32.

10. Chen, Z., Zhu, Y. & Zhu, C. (2016). Adaptive bridge estimation for high-dimensional regression models. Journal of Inequalities and Applications, 1, 258.

11. Endelman, J. B. (2011). Ridge regression and other kernels for genomic selection with R package rrBLUP. The plant genome, 4(3).

12. Eraslan, G., Avsec, Z^., Gagneur, J. & Theis, F.J. (2019). Deep learning: new computational modelling techniques for genomics. Nature Reviews Genetics, 20(7), 389–403.

13. Estaghvirou, S. B. O., Ogutu, J. O., Schulz-Streeck, T., Knaak, C., Ouzunova, M., Gordillo, A., & Piepho, H. P. (2013). Evaluation of approaches for estimating the accuracy of genomic prediction in plant breeding. BMC Genomics, 14(1), 1–21.

14. Fan, J. & Li, R. (2001). Variable selection via nonconcave penalized likelihood and its oracle properties. Journal of the American Statistical Association, 96, 1348–1360.

15. Fan, J. & Peng, H. (2004). Nonconcave penalized likelihood with a diverging number of parameters. Annals of Statistics, 32, 928–961.

16. Frank, I.E. & Friedman, J.H. (1993). A statistical view of some chemometrics regression tools (with discussion). Technometrics, 35, 109–148.

17. Friedman, J. (2001). Greedy function approximation: a gradient boosting machine. Annals of Statistics, 29, 1189–1232.

18. Friedman, J., Hastie, T. & Tibshirani, R. (2010). A note on the group lasso and sparse group lasso. arXiv preprint 1001.0736.

19. Friedman, J., Hastie, T., Tibshirani, R., Narasimhan, B., Tay, K., Simon, N., Qian, J. (2022). Package ‘glmnet’. Journal of Statistical Software. 2010a, 33(1).

20. Fu, W.J. (1998). Penalized regressions: The bridge versus the lasso. Journal of Computational and Graphical Statistics, 7, 397–416.

21. Grandvalet, Y. (1998). Least absolute shrinkage is equivalent to quadratic penalization. International Conference on Artificial Neural Networks, 201–206). Springer, London.

22. Greenwell, B., Boehmke, B., Cunningham, J., Developers, G. B. M. & Greenwell, M. B. (2019). Package ‘gbm’.

23. Hastie, T.J., Tibshirani, R. & Friedman, J. (2009). The elements of statistical learning, Second edition, New York: Springer.

24. Hayes, B. J., Visscher, P. M. & Goddard, M. E. (2009). Increased accuracy of artificial selection by using the realized relationship matrix. Genetics Research, 91(1), 47–60.

25. Heslot, N., Yang, H.P., Sorrells, M.E. & Jannink, J.L. (2012). Genomic selection in plant breeding: a comparison of models. Crop Science, 52, 146–160.

26. Hoerl, A.E. & Kennard, R.W. (1970). Ridge regression: biased estimation for non-orthogonal problems. Technometrics, 12, 55–67.

27. Huang, J., Ma, S., Xie, H. & Zhang, C-H. (2009). A group bridge approach for variable selection. Biometrika, 96, 339–355.

28. Huang, J., Horowitz, J.L. & Ma, S. (2008). Asymptotic properties of bridge estimators in sparse high-dimensional regression models. Annals of Statistics, 36, 587–613.

29. Huang, J. & Zhang, T. (2010). The benefit of group sparsity. Annals of Statistics, 38, 1978–2004.

30. Huang, J., Breheny, P. & Ma, S. (2012). A Selective Review of Group Selection in High-Dimensional Models. Statistical Science, 27(4), 10.1214/12-STS392.

31. Kim, Y., Choi, H. & Oh, H. S. (2008). Smoothly clipped absolute deviation on high dimensions. Journal of the American Statistical Association, 103(484), 1665–1673.

32. Kingma, D.P. & Ba, J.L. (2014). Adam: A method for stochastic optimization. arXiv preprint 1412.6980.

33. Knight, K. & Fu, W. (2000). Asymptotics for Lasso-type estimators. Annals of Statistics, 28, 356–1378.

34. Liaw, A. & Wiener, M. (2002). Classification and regression by randomForest. R News, 2, 18–22.

35. Lim, M. & Hastie, T. (2015). Learning interactions via hierarchical group-lasso regularization. Journal of Computational and Graphical Statistics, 24(3), 627–654.

36. Mazumder, R., Friedman, J.H. & Hastie, T. (2011). Sparsenet: Coordinate descent with nonconvex penalties. Journal of the American Statistical Association, 106(495), 1125–1138.

37. Meyer, D., Dimitriadou, E., Hornik, K., Weingessel, A., Leisch, F., Chang, C. C. et al. (2019). Package ‘e1071’. The R Journal.

38. Meuwissen, T. H., Hayes, B. J. & Goddard, M. (2001). Prediction of total genetic value using genome-wide dense marker maps. Genetics, 157(4), 1819–1829.

39. Min, S., Lee, B. & Yoon, S. (2017). Deep learning in bioinformatics. Briefings in Bioinformatics, 18(5), 851–869.

40. Montesinos-López, A., Montesinos-López, O.A., Gianola, D., Crossa, J. & Hernández-Suárez, C.M. (2018). Multi-environment genomic prediction of plant traits using deep learners with dense architecture. G3: Genes, Genomes, Genetics, 8(12), 3813–3828.

41. Montesinos-López, O.A., Montesinos-López, A., Crossa, J., Gianola, D., Hernández-Suárez, C.M., & Martín-Vallejo, J. (2018). Multi-trait, multi-environment deep learning modeling for genomic-enabled prediction of plant traits. G3: Genes, Genomes, Genetics, 8(12), 3829–3840.

42. Montesinos-López, O.A., Martín-Vallejo, J., Crossa, J., Gianola, D., Hernández-Suárez, C.M., Montesinos-López, A., Philomin J. & Singh, R. (2019). A benchmarking between deep learning, support vector machine and Bayesian threshold best linear unbiased prediction for predicting ordinal traits in plant breeding. G3: Genes, Genomes, Genetics, 9(2), 601–618.

43. Montesinos-López, O.A., Martín-Vallejo, J., Crossa, J., Gianola, D., Hernández-Suárez, C.M., Montesinos-López, A., Juliana, P. & Singh, R., (2019). New deep learning genomic-based prediction model for multiple traits with binary, ordinal, and continuous phenotypes. G3: Genes, Genomes, Genetics, 9(5), 1545–1556.

44. Ogutu, J.O., Piepho, H.P. & Schultz-Streeck, T. (2011). A comparison of random forests, boosting and support vector machines for genomic selection. BMC Proceedings, 5(3), BioMed Central Ltd.

45. Ogutu, J.O., Schulz-Streeck, T. & Piepho H-P. (2012). Genomic selection using regularized linear regression models: ridge regression, lasso, elastic net and their extensions. BMC Proceedings, 6(2), BioMed Central Ltd.

46. Ogutu, J.O. & Piepho, H-P. (2014). Regularized group regression methods for genomic prediction: Bridge, MCP, SCAD, group bridge, group lasso, sparse group lasso, group MCP and group SCAD. BMC Proceedings, 8(5), BioMed Central Ltd.

47. Park, C. & Yoon, Y.J. (2011). Bridge regression: adaptivity and group selection. Journal of Statistical Planning and Inference, 141, 3506–3519.

48. Percival, D. (2011). Theoretical properties of the overlapping groups lasso. Electronic Journal of Statistics, 6, 269–288.

49. Pérez-Enciso, M. & Zingaretti, L.M. (2019). A Guide on Deep Learning for Complex Trait Genomic Prediction. Genes, 10(7), p.553.

50. Piepho H-P. (2009). Ridge regression and extensions for genomewide selection in maize. Crop Science, 49, 1165–1176.

51. Piepho, H-P., Ogutu, J.O., Schulz-Streeck, T., Estaghvirou, B., Gordillo, A. & Technow, F. (2012). Efficient computation of ridge-regression best linear unbiased prediction in genomic selection in plant breeding. Crop Science, 52, 1093–1104.

52. Poignard, B. (2020). Asymptotic theory of the adaptive Sparse Group Lasso. Annals of the Institute of Statistical Mathematics, 72(1), 297–328.

53. Ruder, S. (2016). An overview of gradient descent optimization algorithms. arXiv preprint 1609.04747.

54. Ruppert, D., Wand, M. P., & Carroll, R. J. (2003). Semiparametric regression. Cambridge University Press.

55. Schonlau, M. (2005). Boosted regression (boosting): An introductory tutorial and a Stata plugin. The Stata Journal, 5(3), 330–354.

56. Simon, N., Friedman, J., Hastie, T. & Tibshirani, R. (2013). A sparse-group lasso. Journal of Computational and Graphical Statistics, 22, 231–245.

57. Tibshirani, R. (1996). Regression shrinkage and selection via the lasso. Journal of the Royal Statistical Society, Series B, 58, 267–288.

58. Vapnik, V. (1995). The Nature of Statistical Learning Theory. Springer, New York.

59. Xiao, N. & Xu, Q. S. (2015). Multi-step adaptive elastic-net: reducing false positives in high-dimensional variable selection. Journal of Statistical Computation and Simulation, 85(18), 3755–3765.

60. Xie, L. (2009). Randomly split SAS data set exactly according to a given probability Vector. https://silo.tips/download/randomly-split-sas-data-set-exactly-according-to-a-given-probability-vector

61. Yuan, M. & Lin, Y. (2006). Model selection and estimation in regression with grouped variables. Journal of the Royal Statistical Society, Series B, 68, 49–67.

62. Yue, T. & Wang, H. (2018). Deep learning for genomics: A concise overview. arXiv preprint 1802.00810.

63. Zhang, C-H. (2007). Penalized linear unbiased selection. Department of Statistics and Bioinformatics, Rutgers University, Technical Report #2007-003.

64. Zhang, C.H. & Huang, J. (2008.) The sparsity and bias of the lasso selection in high-dimensional linear regression. The Annals of Statistics, 36, 1567–1594.

65. Zhang, C-H. (2010). Nearly unbiased variable selection under minimax concave penalty. Annals of Statistics, 38, 894–942.

66. Zhou, N. & Zhu, J. (2010). Group variable selection via a hierarchical lasso and its oracle property. Statistics and its Interface, 3, 557–574.

67. Zou, H. & Hastie, T. (2005). Regularization and variable selection via the elastic net. Journal of the Royal Statistical Association, Series B, 67, 301–320.

68. Zou, H. (2006). The adaptive lasso and its oracle properties. Journal of the American Statistical Association, 101, 1418–1429.

69. Zou, H,. Hastie, T. & Tibshirani, R. (2006). Sparse principal component analysis. Journal of Computational and Graphical Statistics, 15(2), 265–286.

70. Zou, H. & Zhang, H.H. (2009). On the adaptive elastic-net with a diverging number of parameters. The Annals of Statistics, 37(4), 1733–1751.

71. Zou, J., Huss, M., Abid, A., Mohammadi, P., Torkamani, A. & Telenti, A. (2019). A primer on deep learning in genomics. Nature Genetics, 51(1), 12–18.

